# Chromosome-scale genome assembly of *Eustoma grandiflorum*, the first complete genome sequence in family Gentianaceae

**DOI:** 10.1101/2021.09.09.459690

**Authors:** Kenta Shirasawa, Ryohei Arimoto, Hideki Hirakawa, Motoyuki Ishimorai, Andrea Ghelfi, Masami Miyasaka, Makoto Endo, Saneyuki Kawabata, Sachiko Isobe

## Abstract

*Eustoma grandiflorum* (Raf.) Shinn., is an annual herbaceous plant native to the southern United States, Mexico, and the Greater Antilles. It has a large flower with a variety of colors and an important flower crop. In this study, we established a chromosome-scale *de novo* assembly of *E. grandiflorum* by integrating four genomic and genetic approaches: (1) Pacific Biosciences (PacBio) Sequel deep sequencing, (2) error correction of the assembly by Illumina short reads, (3) scaffolding by chromatin conformation capture sequencing (Hi-C), and (4) genetic linkage maps derived from an F_2_ mapping population. The 36 pseudomolecules and unplaced 64 scaffolds were created with total length of 1,324.8 Mb. Full-length transcript sequencing was obtained by PacBio Iso-Seq sequencing for gene prediction on the assembled genome, Egra_v1. A total of 36,619 genes were predicted on the genome as high confidence HC) genes. Of the 36,619, 25,936 were annotated functions by ZenAnnotation. Genetic diversity analysis was also performed for nine commercial *E. grandiflorum* varieties bred in Japan, and 254,205 variants were identified. This is the first report of the construction of reference genome sequences in *E. grandiflorum* as well as in the family Gentianaceae.

## Introduction

*Eustoma grandiflorum* (Raf.) Shinn., commonly known as Lisianthus, prairie gentian, or bluebell gentian, is an annual herbaceous plant native to the southern United States, Mexico, and the Greater Antilles.^1,2^ It has a large flower with a variety of colors, such as white, pink, yellow, purple, and purple-edged white.^3^ *E. grandiflorum* is cultivated around the world and has become one of the ten most popular cut flowers.^4^ It is an important flower crop especially in Japan, ranking fourth in production value in 2017 and third in cultivation area in 2018.^5^ Numerous varieties have been bred in the commercial and public sectors^3^ as both selfed and F_1_ hybrids.

Genus *Eustoma*, which belongs to the family Gentianaceae and the tribe Chironiae, is small, comprising only three species: *E. grandiflorum, E. barkleyi* Standely, and *E. exaltatum* (L.) Salisb. Ex Don.^6^ *E. grandiflorum* was previously called *E. russellianum* and is sometimes classified as a subspecies of *E. exaltatum*.^7^ In the NCBI taxonomy database (https://www.ncbi.nlm.nih.gov/taxonomy), *E. grandiflorum* is registered as a heterotypic synonym of *E. exaltatum* subsp. *russellianum* (Taxonomy ID: 52518). *E. grandiflorum* was previously considered an octoploid^8^, but a recent study suggested that *E. grandiflorum* is a diploid, with a chromosome number of 2n = 2X = 72.^9^

The family Gentianaceae consists of six tribes, 99 genera, and approximately 1,736 species.^10^ The family name Gentianaceae is derived from Gentius, an Illyrian king in the time of Ancient Greece, who discovered the medicinal properties of gentian. As indicated by the origin of the family name, several species in the family, such as *Gentiana trifloral* (gentian) and *Swertia japonica*, have been used as medicinal or herbal plants. However, because they have been relatively less used in industry than other plants have been, the species in this family have not been well studied in modern science, especially in the field of genomics. Chloroplast and plastid genome sequences were reported for several species in the genera *Gentiana*^11–15^ and *Pterygocalyx*^16,17^. Transcriptome analyses were also reported for *G. straminea*^18^, *G. rigescens*^19^, and *Swertia nussotii*^20^. However, as far as we currently know, no whole genome sequencing has been reported on a chromosome scale in a Gentianaceae species.

In this study, we established a chromosome-scale *de novo* assembly of *E. grandiflorum* by integrating four genomic and genetic approaches: (1) Pacific Biosciences (PacBio) Sequel deep sequencing, (2) error correction of the assembly by Illumina short reads, (3) scaffolding by chromatin conformation capture sequencing (Hi-C), and (4) genetic linkage maps derived from an F_2_ mapping population. Full-length transcript sequencing was obtained by PacBio Iso-Seq sequencing for gene prediction on the assembled genome, Egra_v1. Genetic diversity analysis was also performed for nine commercial *E. grandiflorum* varieties bred in Japan. This is the first report of the construction of reference genome sequences in *E. grandiflorum* as well as in the family Gentianaceae. We expect the assembled genome will contribute to the advance of research and breeding in *E. grandiflorum* and will help to identify genes in the family Gentianaceae that will be useful in medicinal and other industries.

## Materials and Methods

### Whole genome sequencing and assembly

An *E. grandiflora* inbred line, 10B-620, bred at Nagano Vegetable and Ornamental Crops Experimental Station, was used for whole genome sequencing with Illumina short reads and PacBio long reads. The Genomic DNA was extracted from young leaves with the use of the Genomic DNA Extraction Column (Favorgen Biotech Corp., Ping-Tung, Taiwan) for short reads and the Genomic-tips Kit (QIAGEN, Germantown, MD, USA) for long reads.

An Illumina paired-end (PE) library was constructed with an expected insert size of 500 bp. Library sequencing was performed by an Illumina HiSeq (Illumina, San Diego, CA, USA) system with a read length of 101 nt (Supplementary Table S1). A genome size of 10B-620 was estimated based on kmer-frequency analysis with short reads by using Jellyfish ver. 2.1.1^21^.

A long-read sequence library was prepared using the SMRTbell Express Template Prep Kit 1.0 (PacBio, Menlo Park, CA, USA). The size selection of the library was performed by BluePippin (Sage Science, Beverly, MA, USA) to remove DNA fragments less than 15 kb in length, and the library was then sequenced using the Sequel system (PacBio) with 14 SMRT cells.

The sequence reads were assembled using FALCON Unzip v.1.8.1^22^ with default parameters, and the generated primary contig sequences were polished twice using ARROW ver. 2.2.1 implemented in SMRT Link v.5.0 (PacBio). Illumina PE reads were then used for further error correction of the contig sequences by using Pilon 1.22^23^.

### Linkage map construction

An F_2_ mapping population named 10B-58 was developed from reciprocal crosses between 10B-620 and an inbred *E. grandiflorum* line, 10B-503. The number of F_2_ individuals used for linkage map construction was 104. Variants (SNPs and Indels) segregating in the F_2_ population were detected by sequencing the dd-RAD-Seq and GRAS-Di libraries. Library construction was performed according to Shirasawa et al. (2016)^24^ for dd-RAD-Seq and to Miki et al. (2020)^25^ for GRAS-Di. Both libraries were sequenced using Illumina Hiseq 2000 (Illumina). A variant call was performed by bcftools 0.1.19 mpileup in Samtools^26^.

A linkage map was constructed twice by using Lep-MAP3^27^ and MSTmap^28^. The map created using Lep-MAP3 (hereinafter Lep map) was constructed with the variants identified on the FALCON-unzip contigs and was used to split misassembled contig sequences by comparing the SNP positions on the contigs and with those on the linkage groups. The default parameters were used in the Lep-Map3, and the male map positions are shown in this study. The map created using MSTmap (hereinafter MST map) was constructed for the revision of the chromosome-scale scaffolds after the Hi-C analysis. The following parameters were used to construct the linkage map: distance_function = kosambi, cut_off_p_value = 1e-12, no_map_distance = 20, no_map_size = 2, missing_threshold = 0.2, estimation_before_clustering = yes, detect_bad_data = no, objective_function = COUNT.

### Hi-C scaffolding and construction of chromosome-level scaffolds

A Hi-C library was constructed from young leaves of 10B-620 using a Proximo Hi-C Plant Kit (Phase Genomics, Seattle, WA, USA). The library was sequenced by Illumina NextSeq500, and the obtained PE reads were aligned onto the scaffolds by BWA^29^. Chromosome-scale scaffolds were created by using the Proximo Hi-C genome scaffolding platform (Phase Genomics) in a method similar to that described by Bickhart et al. (2017)^30^. Juicebox^31^ was then used to correct scaffolding errors. The Hi-C scaffolds were then cut and reordered by using ALLMAPS^32^ and Ragoo^33,34^ with the MST map as a reference, and the chromosome-level scaffold sequences were determined.

Assembly quality was assessed by benchmarking universal single-copy orthologs (BUSCOs) sequences using BUSCO v3.0.^34^ Repetitive sequences in the assembled genome were identified by RepeatMasker 4.0.7 (http://www.repeatmasker.org/RMDownload.html) for known repetitive sequences registered in Repbase (https://www.girinst.org/repbase/) and de novo repetitive sequences defined by RepeatModeler 1.0.11 (http://www.repeatmasker.org/RepeatModeler).

### Transcriptome sequencing and gene prediction

Total RNAs were extracted from young leaves and buds of 10B-620 by using the RNeasy Plant Mini Kit RNA (QIAGEN). Iso-Seq libraries were created for leaves and buds in accordance with the manufacturer’s protocol (PacBio) and sequenced by a Sequel system with two SMRT cells. The obtained reads were clustered using the Iso-Seq 2 pipeline implemented in SMRT Link ver.5.1.0 (PacBio). The high-quality (hq) Iso-seq sequences were then mapped onto the assembled genome with Minimap2^35^ and collapsed to obtain nonredundant isoform sequences using a module in Cupcake ToFU (https://github.com/Magdoll/cDNA_Cupcake). ORF (open reading frame) prediction on the collapsed sequences was performed using ANGEL (https://github.com/PacificBiosciences/ANGEL). Redundant sequences were then removed by the CD-HIT program,^36^ and nonredundant complete confidence (cc) sequences were mapped onto the assembled genome sequences by GMAP ver. 2020.06.01.^37^

Meanwhile, empirical gene prediction was performed for the repeat masked assembled genome sequences by BRAKER2^38^ with published *E. grandifolum* transcript sequences (Supplementary Table S2). After removal of the redundant variant sequences, the gene sequences predicted by BRAKER v2 were merged with those mapped with cc Iso-Seq sequences. When gene sequences were predicted by both BRAKER v2 and Iso-Seq, the longest CDSs were selected.

In order to classify the predicted gene sequences based on the evidence level, a similarity search was performed against the NCBI NR protein database (http://www.ncbi.nlm.nih.gov) and UniProtKB (https://www.uniprot.org) using DIAMOND^39^ with 60% ≤ similarity, 50% ≤ mapped length ≤ 150%, and E-value ≤ 1E-80. BLASTP searches were also performed for the gene sequences of *Vitis vinifera* (12X)^40^ and *Arabidopsis thaliana* (Araport 11)^41^ with 30% ≤ similarity (*V. vifinera*) or 25% ≤ similarity (*A. thaliana*), 50% ≤ mapped length ≤ 150%, and E-value ≤ 1E-80. Domains were searched by HAMMER v3.3.2 (http://hmmer.org/) with E-value ≤ 1E-30, and TPM values were calculated by Salmon (Ref) with the RNA-Seq reads listed in Supplementary Table S2. The high confidence (HC) gene sequences were selected under the following conditions: TPM value > 0.2, identified protein domain sequences, gene sequence hits in UniProtKB or NR protein database, or *V. vinifera* genes. Transposon elements (TEs) were classified based on the results of similarity searches against UniProtKB. The gene sequences not classified as HC or TE were classified as LC (low confidence). Functional gene annotation was also performed by using a modified version of Hayai annotation,^42^ called ZenAnnotation (https://github.com/aghelfi/ZenAnnotation), in which it was incorporated OrthoDB sequences in order to allow a contaminant detection. The parameters for sequence alignment, performed by diamond with a more-sensitive algorithm, were sequence identity 50%, query cover 50% and subject cover 50%.

### Diversity analysis in nine commercial varieties

Genetic diversity was investigated in nine commercial varieties bred in Japan: Yukitemari, Borelo white, Paleo pink, Exe lavender, La folia, Umi honoka, Robera clear pink, Korezo light pink, and Celeb pink. Plant materials were grown in the field at Takii Co. Ltd. (Shiga, Japan), and the genomic DNA of each was extracted from young leaves with the use of the Genomic DNA Extraction Column (Favorgen Biotech Corp). Whole genome shotgun (WGS) sequencing was performed by using Illumina Hisex X (Illumina) with 150 PE reads. The WGS reads were mapped onto the assembled genome sequences by using Bowtie 2,^43^ and base variants were identified using bcftools 0.1.19 mpileup in Samtools.^29^ Genetic distances were calculated by using the Distance Matrix function in TASSEL 5.^44^ A NJ phylogenetic tree was constructed using MEGA ver 10.1.8.^45^

## Results and Discussion

### Estimation of 10B-620 genome size

WGS reads were obtained for 10B-620 with a total length of 64.26 Gb reads (Supplementary Table S1). The distribution of distinct k-mers (k=17) shows a single large peak at multiplicities of 34, suggesting that 10B-620 was a highly homozygous material (Fig. 1). Based on the identified peak, the genome size of 10B-620 was estimated to be 1,587.6 Mb. Lindsay et al. (1994)^46^ reported that the nuclear DNA content of diploid *E. grandiflorum* was 3.26 ± 0.10 pg DNA per 2C, which suggested that the total base length of the genome was 1512.64 Mb per C. Our genome size estimation was almost similar with that of the previous report.

**Fig. 1.**
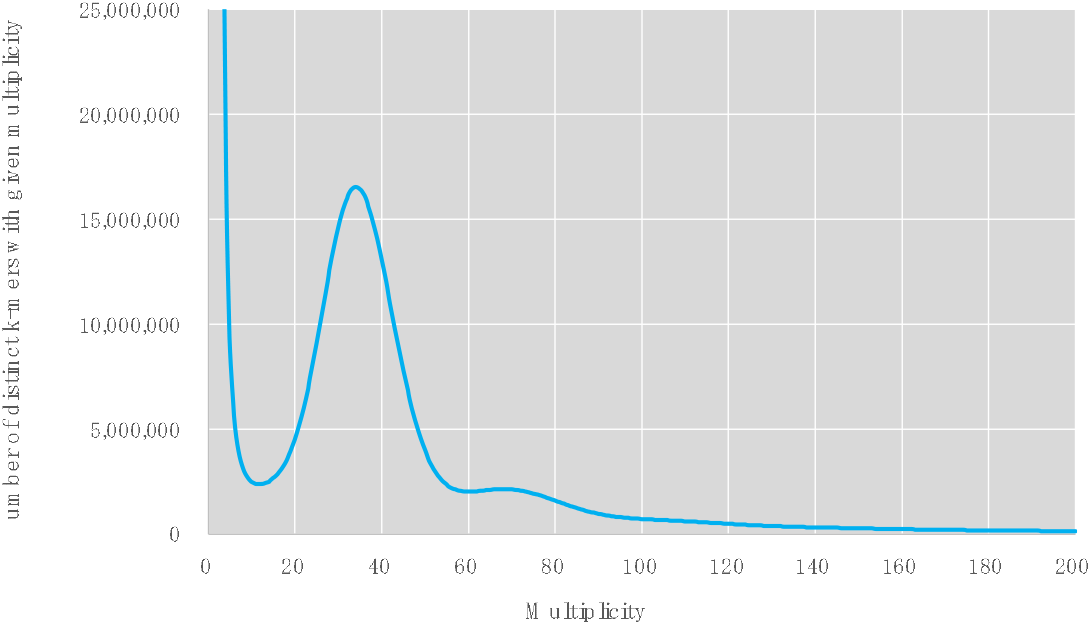
Genome size estimation using Jellyfish with the distribution of the number of distinct kmers (kmer =17) and the given multiplicity values.

### PacBio assembly

A total length of 104.47 Gb of PacBio reads was generated from 14 SMRT cells. The obtained read coverage against the 10B-620 genome was 65.8x. The Falcon unzip assembly generated 675 primary and 7,724 haplotig contigs (Supplementary Table S3). The total length of the primary contigs was 1,322.4 Mb, with an N50 length of 4.70 Mb. The primary contig sequences were then polished with Sequel reads using Arrow, followed by further polishing with the Illumina reads using Pilon. The resultant number of primary sequences was 753, with a total length of 1,324.5 Mb.

### Linkage map construction for identification of misassembly of the PacBio contigs

dd-RAD-Seq and GRAS-Di sequences were obtained for the 104 F_2_ population (10B-58) derived from crosses between 10B-620 and 10B-503. The reads were mapped onto the 753 primary sequences, resulting in the identification of 20,401 (dd-RAD-Seq) and 5,488 (GRAS-Di) base variants. The variants identified from the two libraries were then merged, and a linkage map was constructed by Lep-MAP3. A total of 20 linkage groups (LGs) were generated, with a total length of 2,331.5 cM (Supplementary Fig. 1). The numbers of mapped loci and bins (unique positions of loci) were 17,872 and 1,358, respectively.

A total of 79 contigs were identified as possible misassemblies by comparing the SNP positions between the Lep map and F_2_ genotype segregation patterns. The 79 contigs were split at the points of possible misassembly, and the resultant 753 primary contigs were used for subsequent Hi-C scaffolding (Supplementary Table S3).

### Chromosome-level scaffolding with Hi-C reads and a linkage map

Several different chromosome numbers and ploidies of *E. grandiflorum* have been reported. For example, Rork et al. (1949)^8^ described *E. grandiflorum* as an octoploid, with a chromosome number of 2n = 8X = 72 based on observation of chromosomes in root chips. Griesbach and Bhat (1990)^47^ reported that the basic chromosome number of *E. grandiflorum* was 18 according to a chromosome observation in meiotic metaphase I of diploid *E. grandiflorum.* Meanwhile, Kawakatsu et al. (2021)^9^ suggested that the chromosome number of *E. grandiflorum* was considered to be 2n = 2x = 72, based on the result of SSR linkage map construction and the *E. exaltatum* chromosome number of 2n = 2x = 72 reported by Barba-Gonzalez et al. (2015)^6^. Hence, we constructed chromosome-level scaffolds under the conclusion that the basic chromosome number of *E. grandiflorum* was n = 36.

A total of 589.9 M Hi-C reads were generated and used for scaffolding of the 753 primary contigs with N100. The generated number of scaffolds was 100, including 36 chromosome-level scaffolds. The total length of the 100 scaffolds was 1,324.8 Mb, and the 36 chromosome-level scaffolds occupied 98.8% of the total length.

In order to identify possible misassemblies on the Hi-C scaffolds, a linkage map was reconstructed by using the MST map. The dd-RAD-Seq and GRAS-Di sequences of the F_2_ population were mapped onto the 100 HI-C scaffold sequences. The variants were filtered out with DP ≥ 10 and GQ ≥ 50, and 6,430 variants on the 36 chromosome-level scaffolds were mapped onto 43 LGs. The larger number of LGs than scaffolds suggested possible misassemblies in the process of Hi-C scaffolding.

We then tried to realign the primary contigs that were scaffolded on the wrong position by Hi-C analysis, by referring to base variant positions on the MST map. First, the positions corresponding to the 6,430 variants on the MST map were determined to be against for the primary contigs. As a result, the positions of the 6,430 variants were determined on 360 of the 753 primary contigs. The 360 primary contigs were then aligned on the MST map by using ALLMAPS (here, let’s call the resultant sequences ALLMAPS scaffolds). The 36 chromosome-scale Hi-C scaffold sequences were aligned onto the ALLMAPS scaffolds by using Ragoo with the chimera cut option. After making several manual minor revisions, we created 36 pseudomolecules out of the 36 chromosome-scale scaffolds. The correspondence positions on the pseudomolecules and MST map, as well as the summary of the MST map, are shown in Fig. 4 and Supplementary Tables S4 and S5, respectively.

**Fig. 4.**
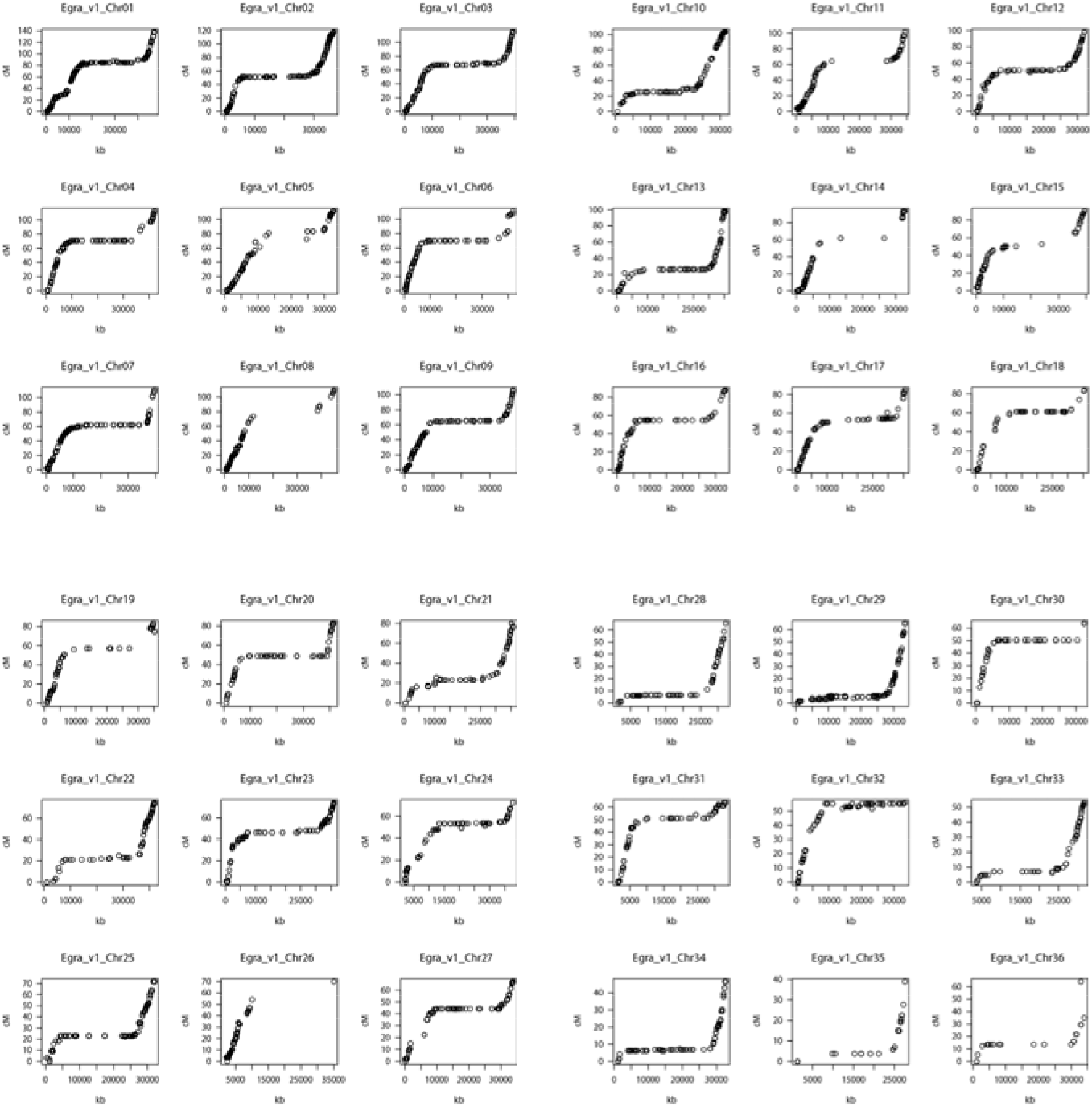
Correlation between physical and genetic distance on the *Eustoma* genome. The genetic distances were calculated from the 10B-581 F_2_ linkage map.

**Fig. 4.**
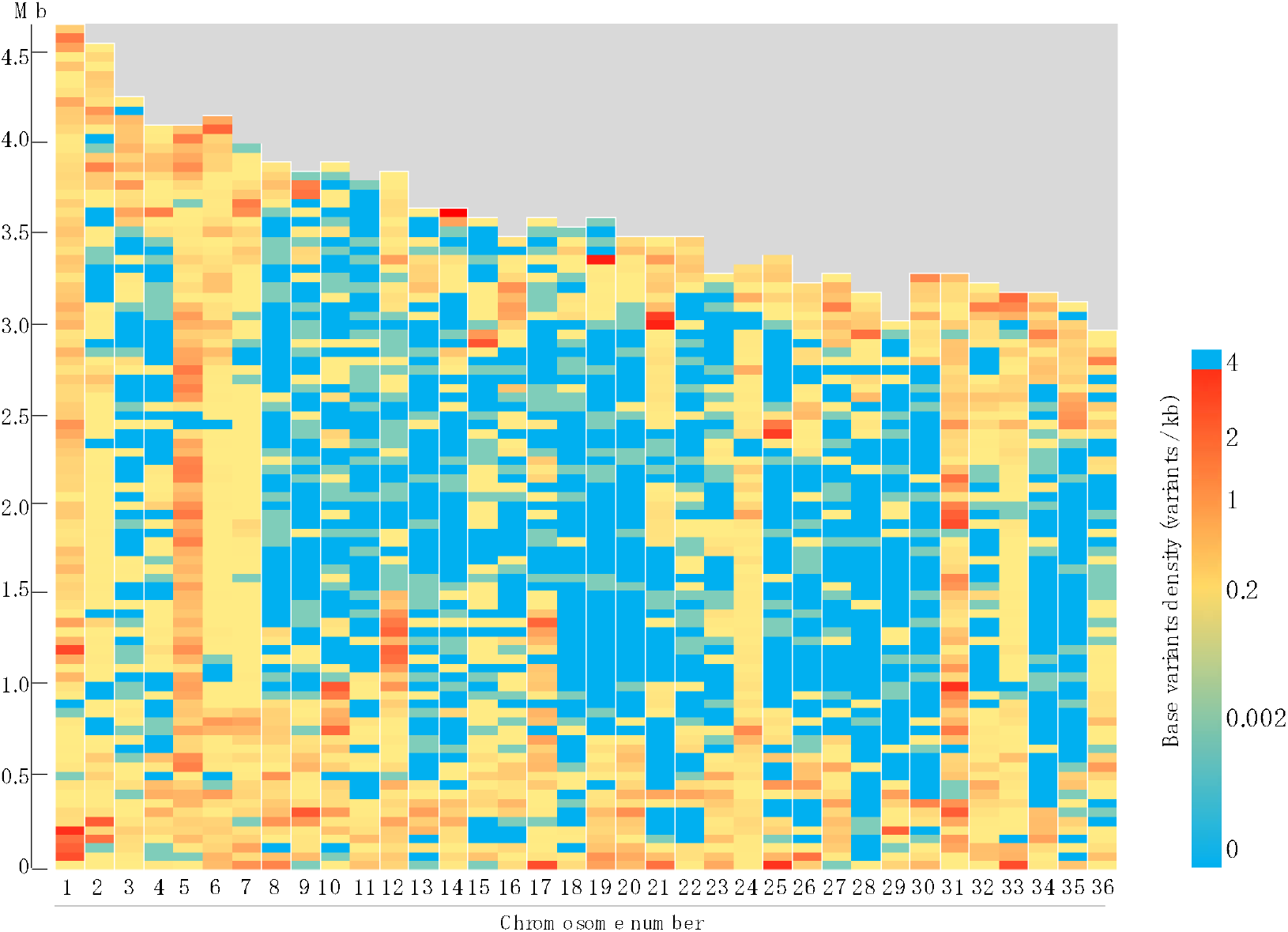
Base variant density on the *Eustoma* genome. The variants were identified with the nine *Eustoma* varieties, and the density (variants/kb) was calculated in each 500 kb windows.

The 36 pseudomolecules and 64 unplaced scaffolds were designated ‘Egra_v1’. The total length of Egra_v1 is 1,324.8 Mb, with a gap length of 329,560 bp (Table 1, Supplementary Table S3). The total length of the 36 pseudomolecules is 1,308.6 Mb, occupying 98.8% of the assembled sequences. The corresponding chromosome numbers of the pseudomolecules are given in longer sequence order. The lengths of the 36 pseudomolecules ranged from 47.12 Mb (Chr01) to 29.6 Mb (Chr36, Supplementary Table S6). When the total lengths are compared with the estimated genome size of 10B-620 (1,587.6 Mb), all of the assembled scaffolds and the 36 pseudomolecules in Egra_v1 cover 83.4% and 82.4% of the genome, respectively.

**Table 1.**
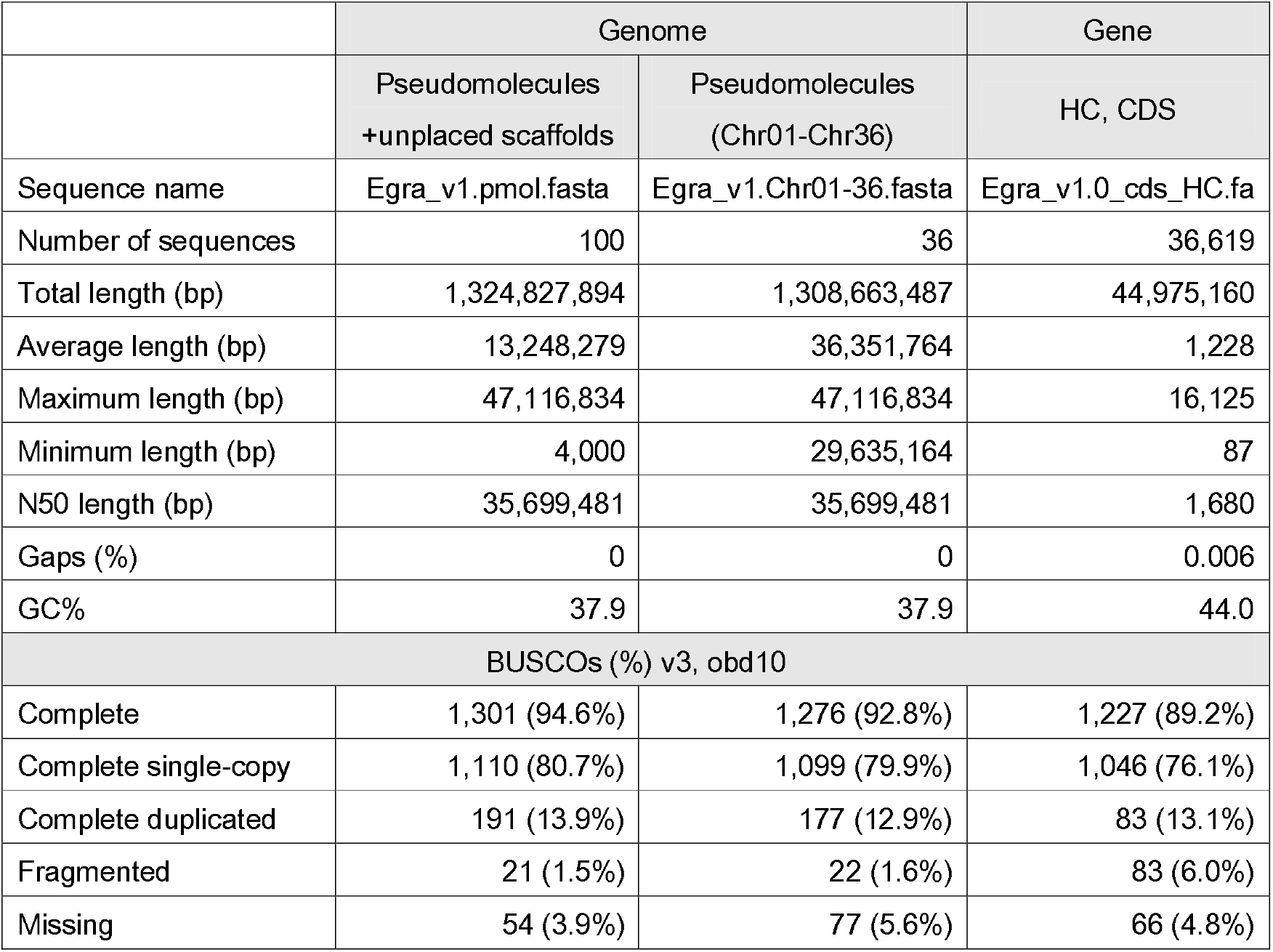
Statistics on the assembled *Eustoma* genome sequences and CDSs (Eqra v1).

The assembly quality of Egra_v1 was investigated by mapping the sequences onto 1,375 BUSCOs (Table 1). The results demonstrated that the number of complete BUSCOs was 1,301 (94.6%), including 1,110 (80.7%) single-copy genes and 191 (13.8%) duplicated genes. There were 21 and 54 fragmented and missing BUSCOs, respectively.

### Gene prediction and annotation

Iso-Seq sequences totaling 735.8 Mb and 982.2 Mb in length were obtained from leaves and young buds, respectively (Supplementary Table S1). The sequences from the two organs were integrated and clustered by Iso-Seq2, and the 50,934 high-quality (hq) sequences were assembled (Supplementary Table S7). The 50,934 sequences were collapsed and filtered based on quality. The resultant 29,132 sequences were then predicted ORF, and 11,175 nonduplicate full-length cDNA sequences were determined, with a total length of 14.0 Mb.

Meanwhile, *de novo* gene prediction was performed on the Egra_v1 genome sequences by using BRAKER2 with the *E. grandiflorum* transcript sequences listed in Supplementary Table S2. As a result, 202,561 candidate genes were predicted on the genome, with a total length of 242 Mb. The predicted gene sequences were merged with the 11,175 full-length cDNA sequences, and the resultant 200,998 sequences were classified as HC, LC, or TE based on evidence level.

The numbers of predicted gene sequences classified as HC, LC, and TE were 36,619, 76,014, and 88,365, respectively (Supplementary Table S8, Table 1). The percentage of complete BUSCOs in HC was 89.2%, while those in LC and TE were 1.2% and 1.5%, respectively. Therefore, most of the protein coding gene sequences were designated as HC.

Functional gene annotation was performed by using a modified version of Hayai annotation with refereeing through the Kusaki database (http://pgdbjsnp.kazusa.or.jp/app/kusakidb). The numbers of functional annotated genes in HC, LC and TE were 25,936, 16,929 and 54,565, respectively (Supplementary Table S9). The most frequently listed species as top hit species against the *E. grandiflorum* genes were *Coffea arabica* (40.3%), then *C. canephora* (9.1%) and *C. eugenioides* (3.4%). The frequently observed top hit families were Rubiaceae, Solanaceae and Nyssaceae, occupying 53.1%, 8.0% and 5.5% of the top hit family list, respectively (Supplementary Table S10). There were 10,356 genes annotated with GO and GOSLIM-PIR terms, 13,205 with PFAM, 13,826 with InterPro and 1,467 with EC (Supplementary table S11-S15).

### Diversity analysis in nine commercial varieties

Illumina PE reads of the nine *E. grandiflorum* varieties bred by Japanese commercial companies were mapped onto Egra_v1 to detect base variants. A total of 16,412,137 candidate variants were identified and filtered according to the following conditions: QUAL 250 ≤, DP 10 ≤, GQ 10 ≤, max-missing = 0.8, and excluding MAF (minor allele frequency) = 0 or 0.5. The remaining number of variants was 254,205. The base variant density on Egra_v1 is shown in Fig. 4. In most of the chromosomes, fewer variants were observed in the middle. However, a few chromosomes, such as Chr5 and Chr31, showed more variants in the middle. Phylogenetic analysis showed that ‘Borelo white’ was genetically distant from the other eight varieties (Fig. 5).

**Fig. 5.**
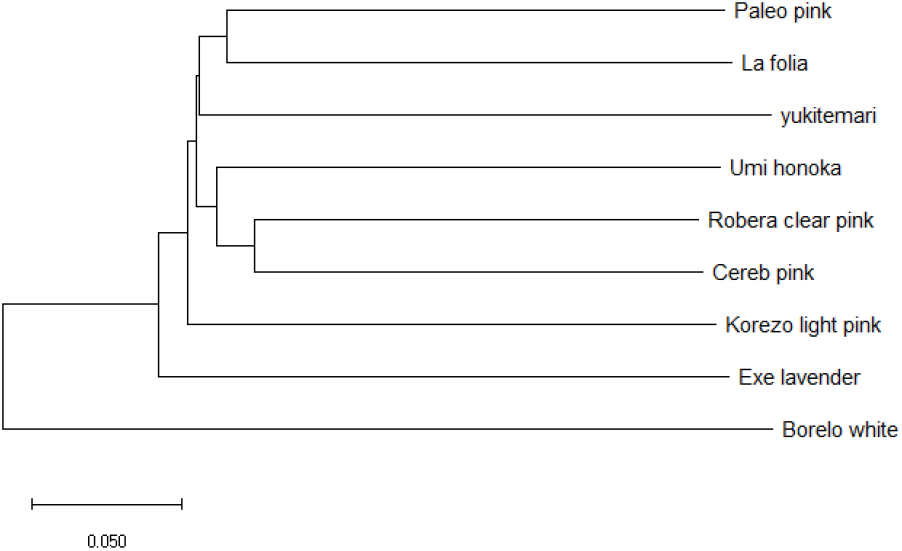
Phylogenetic tree of the nine *Eustoma* varieties based on 254,205 variants.

## Conclusion

In this study we established a chromosome-scale genome assembly of *E. grandiflorum*, the first complete genome sequence in the family Gentianaceae. The assembled genome covered 83.4% of the estimated genome, and the 36 pseudomolecules occupied 98.8% of the assembled genome. In addition, a total of 36,619 protein coding genes were identified on the assembled genome with high confidence. The resultant genome assembly will be useful for genetic and genomic studies and will deepen our understanding of the species in the genus *Eustoma* and the family Gentianaceae.

## Data availability

The assembled genome sequences have been submitted to the DDBJ/ENA/NCBI public sequence databases under the BioProject ID PRJDB12119. The assembled genome and gene sequences, the SNPs of the nine commercial varieties, and the MST map information are available at Plant GARDEN (https://plantgarden.jp/en/list/t52518).

## Acknowledgments

We acknowledge technical assistance by Akiko Watanabe, Yoshie Kishida, Shinobu Nakayama, Shigemi Sasamoto, Hisano Tsuruoka, Chiharu Minami, Mitsuyo Kohara, Takaharu Kimura, Manabu Yamada, Tshunakazu Fujishiro, Akiko Komaki, Akiko Obara, Rie Aomiya, and Taeko Shibazaki of the Kazusa DNA Research Institute. This work was supported by Research Funds provided by Takii Co. Ltd. and the Kazusa DNA Research Institute.

---

**Supplementary Fig. S1.**
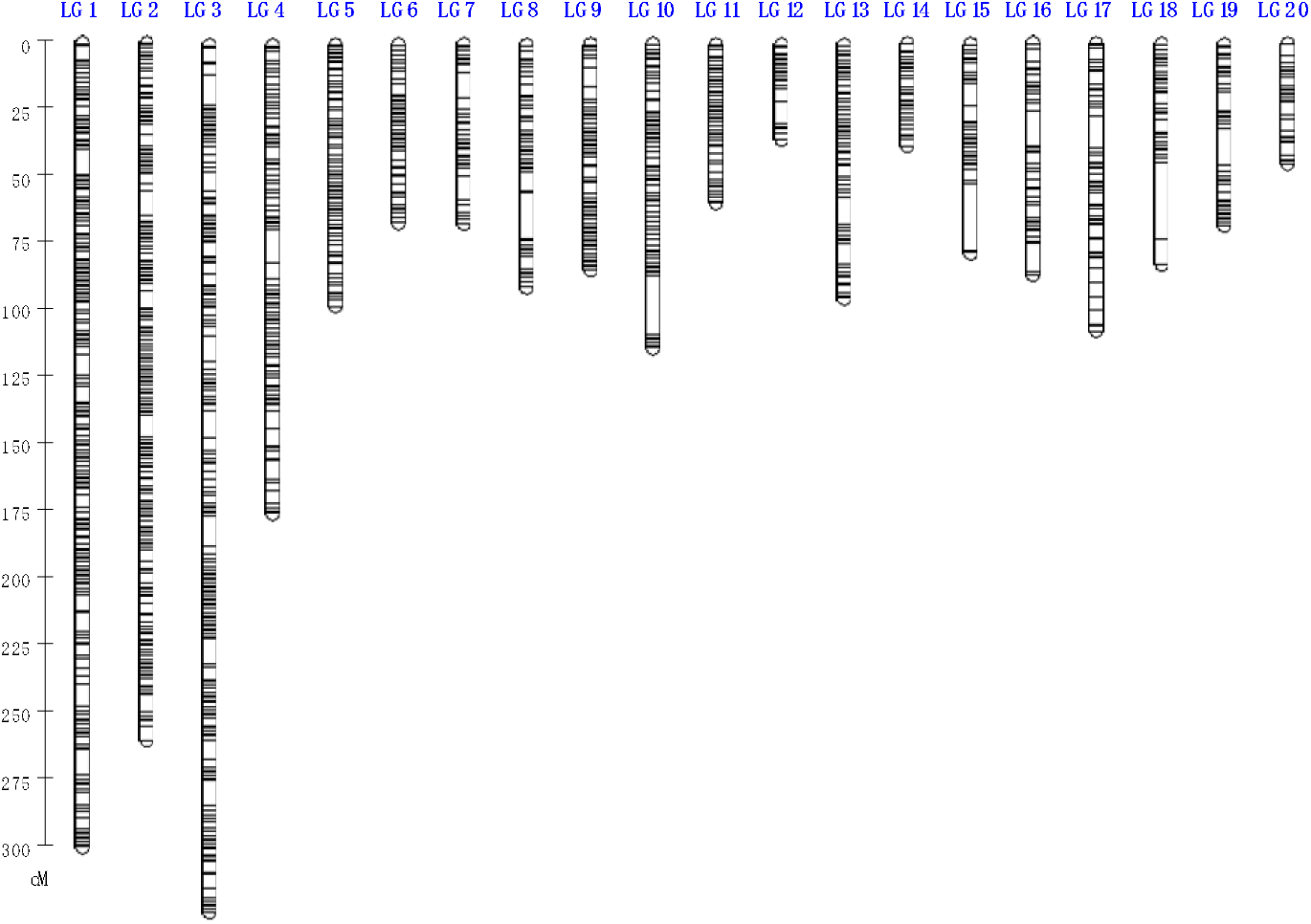
Lep-MAP3 linkage map (male) of the 10B-58 F_2_ mapping population derived from crosses between 10B-620 and 10B-503. The numbers of mapped loci and bins were 17,873 and 1,358, respectively.

